# Simplified high-throughput methods for deep proteome analysis on the timsTOF Pro

**DOI:** 10.1101/657908

**Authors:** Jarrod J Sandow, Giuseppe Infusini, Laura F Dagley, Rune Larsen, Andrew I Webb

## Abstract

Recent advances in mass spectrometry technology have seen remarkable increases in proteomic sequencing speed, while improvements to dynamic range have remained limited. An exemplar of this is the new timsTOF Pro instrument, which thanks to its trapped ion mobility, pushes effective fragmentation rates beyond 100Hz and provides accurate CCS values as well as impressive sensitivity. Established data dependent methodologies underutilize these advances by relying on long analytical columns and extended LC gradients to achieve comprehensive proteome coverage from biological samples. Here we describe the implementation of methods for short packed emitter columns that fully utilize instrument speed and CCS values by combining rapid generation of deep peptide libraries with enhanced matching of single shot data dependent sample analysis. Impressively, with only a 17 minute separation gradient (50 samples per day), the combination of high performance chromatography and CCS enhanced library based matching resulted in an average of 5,931 protein identifications within individual samples, and 7,244 proteins cumulatively across replicates from HeLa cell tryptic digests. Additionally, an ultra-high throughput setup utilizing 5 min gradients (180 samples per day) yielded an average of 3,666 protein identifications within individual samples and 4,659 proteins cumulatively across replicates. These workflows are simple to implement on available technology and do not require complex software solutions or custom-made consumables to achieve high throughput and deep proteome analysis from biological samples.

## Introduction

The orchestration of essentially all biological processes is accomplished by proteins, the actors that create and control any given phenotype. The large-scale identification and quantification of proteins – termed proteomics – has facilitated researchers in studying and understanding complex phenotypes, encompassing a systems-wide view of the deep complexity of living systems. Further development in proteomics approaches hold exciting promise for advancing our understanding of cell biology and biological systems in health and disease. Mass spectrometry (MS)-based approaches to proteomics has become the technology of choice for systems-wide analysis of proteins, their post-translational regulation and interactions. Over the past few decades, technology and methodological improvements have pushed proteomics to the forefront of addressing complex questions in cell biology and biomedicine. These improvements have been driven by advances in chromatography, informatics and primarily, developments that increase the accuracy, sensitivity and speed of MS instruments.

Proteomics approaches predominantly consist of a ‘bottom-up’ workflow, where proteins are extracted from a sample of interest and enzymatically hydrolyzed to create shorter peptides that are more amenable to high resolution chromatography and mass spectrometric analysis. The resulting highly complex mixture of peptides are separated via nano-flow ultra-high performance liquid chromatography (UHPLC) and introduced into the mass spectrometer by electrospray ionization. Operating in ‘data dependent’ mode of acquisition, mass spectrometers detect suitable peptide precursor ions (MS) and subjects them to higher energy collisions to induce fragmentation (MS/MS). High precision mass spectrometers detect in excess of 100,000 molecular features^1^, of which, only a small proportion are identified. Improvements to shotgun proteomics approaches to increase the number of identifications in any given sample, are hampered by increases in multiplexed (chimeric) spectra. Thus, any improvements in sensitivity, dynamic range or chromatographic peak capacity all increase the number of co-eluting peptides, providing diminishing returns using traditional approaches.

Many MS analyzers have been employed in shotgun proteomics, however, time-of-flight (TOF) instruments in particular have properties ideal for the analysis of complex peptide mixtures. TOF instrument performance has steadily improved over the years providing resolving power >35,000 within <100μs of acquisition time^2^. The very fast acquisition rates of TOF instruments also permits coupling to fast orthogonal separation techniques such as ion-mobility spectrometry (IMS)^3^. IMS separates ions in the gas phase based in their collisional cross-sectional area (CCS, Ω), providing information about the size and shape of the molecules. IMS typically has separation times of 10-100 milliseconds. As the separation speed of IMS (milliseconds) sits between that of liquid chromatography (seconds) and MS TOF detection (micro seconds), the nesting of IMS between LC and MS provides an additional dimension of orthogonal information, drastically increasing the effective separation of co-eluting and near isobaric features. However, traditional IMS approaches such as drift tubes, have proved challenging due to device sizes, voltages required and limits to the proportion of the incoming ion-beam that can be utilized^4^.

Trapped Ion Mobility (TIMS) provides a fundamentally new approach to IMS, using an electrostatic gradient tunnel to hold ions against the incoming gas stream. A molecules position within the TIMS device is a function of its CCS area, thus providing high resolution IMS in a short space at low voltages and has the additional benefit of concentrating ions from the incoming ion beam to increase sensitivity. The recent implementation of tandem, sequential accumulation and separation TIMS tunnels now allows for 100% duty cycle to be achieved^5^. The recently implemented ‘Parallel Accumulation – SErial Fragmentation’ (PASEF) method implemented on the timsTOF Pro takes further advantage of the ion concentration and IMS separation, staggering fragmentation with the near linear correlation of peptide mz and CCS (where low mz = low mobility and high mz = high mobility), improving the effective rate of precursor fragmentation beyond 100Hz^6^.

Whilst improvements in MS sequencing speed provide more complete datasets, the overall performance and capability of LC-MS system still relies heavily on chromatographic separation performance. In the pursuit of ever-increasing depths of coverage, columns and separation gradients have tended to become longer. However, as MS dynamic range is one of the main limitations, these approaches produce diminishing returns, resulting in modest increases in proteome coverage at the expense of very large amounts of MS time and consequently low through-put. The release of instruments with drastically improved sequencing speeds opens up new possibilities that can potentially provide much greater depth of information per unit of MS time^7^, which will be of particular importance in the adoption of LC-MS into clinical utility. Thus, strategies that combine orthogonal pre-fractionation to increase the effective dynamic range, run on ultra-short gradients, is an effective method for overcoming the dynamic range limitation of MS instruments^8^.

Here we present an optimized and easily implementable approach for generating deep proteomes in very short timescales enabling increased through-put. The method combines high pH reversed phase stage-tip fractionation for peptide feature library generation on short gradients on high peak capacity commercially available columns to take full advantage of the timsTOF Pro’s >100 Hz tandem MS scanning speed. Additionally, accurate CCS values are exploited^9^ to provide deep library-based matching of individual samples running on short gradients for drastically improved utilization of MS analysis time and increasing peptide and protein identifications.

## Materials and Methods

### Hela tryptic digest

Hela cell tryptic peptides were obtained from the commercially prepared Pierce HeLa Protein Digest Standard (Thermo Fisher). Each vial was reconstituted in 2% acetonitrile (ACN)/1% formic acid (FA) in MilliQ water to a concentration of 200ng/μl before being aliquoted and frozen at −80°C prior to analysis.

### Plasma collection and digestion

Blood was collected into tubes containing EDTA and centrifuged for 10 min at 1800 g, supernatant transferred to a new tube and centrifuged again for 15 min at 2000 g to harvest plasma. Blood was sampled from a healthy donor, who provided written informed consent, with prior approval of the ethics committee of the Walter and Eliza Hall Institute. The sample was digested using the SP3 protocol as described by Hughes et al.^10^ with some modifications. A 1:1 combination mix of two of magnetic carboxylate beads was used (Sera-Mag Speed beads, #45152105050250, #65152105050250, GE Healthcare). Beads were prepared fresh by rinsing with water three times prior to use at a stock concentration of 20 μg/μL. The plasma sample (2.5μl) was simultaneously reduced and alkylated in a buffer containing 10 mM Tris HCl pH 7.4/10 mM Tris (2-Carboxyethyl) phosphine (TCEP)/5.5 mM 2-chloracetamide (2-CAA) by heating at 95°C for 10 mins. Carboxylate beads (4 μl) were added to the sample with ACN (70% final concentration v/v) and incubated at RT for 18 mins. Samples were placed on a magnetic rack (Ambion, Thermo Fisher Scientific), supernatant discarded, and the beads washed twice with 70% ethanol and once with neat ACN. ACN was completely evaporated from the tube using a CentriVap (Labconco) prior to the addition of 80μl digestion buffer (10% 2,2,2-Trifluoroethanol (TFE)/100 mM NH_4_HCO_3_) containing 4μg Trypsin-gold (Promega, V5280) and 4μg Lys-C (Wako, 129-02541). Enzymatic digestion proceeded for 1 hr at 37 °C using the ThermoMixer C (Eppendorf) shaking at 400 rpm. Following the digest, sample was placed on a magnetic rack and the supernatant containing peptides was collected and an additional elution (50μl) was performed using 2% dimethyl sulfoxide (DMSO, Sigma) prior sonication in a water bath for 1 min. Sample was lyophilised to dryness using a CentriVap (Labconco) before reconstitution in 250μl 2% ACN, 1% FA and frozen in 3 aliquots prior to analysis.

### High-pH fractionation

50μg of peptides from either digested plasma or 20μg of a Hela tryptic digest (Pierce, Thermo Fisher) were resuspended in 10mM Ammonium Formate pH 10. Peptides were separated into 12 fractions using a stage-tip containing 4 X C18 plugs. The stage-tips were activated using isopropanol, washed with 60% ACN in 10mM Ammonium Formate pH 10 and re-equilibrated using 10mM Ammonium Formate pH 10. Samples were then loaded onto the stage-tips, washed twice using 10mM Ammonium Formate pH 10 and eluted into fractions using an escalating concentration of ACN in 10mM Ammonium Formate pH 10 (2.75, 3.75, 5, 6, 7, 8, 9, 10, 13, 17.5, 25, 60% ACN). Fractions were lyophilised to dryness using a CentriVap (Labconco) before reconstitution in 2% ACN, 1% FA prior to analysis.

### 11 and 50 samples per day UHPLC settings

Samples were analyzed on a nanoElute (plug-in V1.1.0.27; Bruker, Germany) coupled to a timsTOF Pro (Bruker) equipped with a CaptiveSpray source. Peptides were separated on a 15cm X 75μm analytical column, 1.6μm C18 beads with a packed emitter tip (IonOpticks, Australia). The column temperature was maintained at 50°C using an integrated column oven (Sonation GmbH, Germany). The column was equilibrated using 4 column volumes before loading sample in 100% buffer A (99.9% MilliQ water, 0.1% FA) (Both steps performed at 980bar). For the 11 samples per day method, samples were separated at 400nl/min using a linear gradient from 2% to 25% buffer B (99.9% ACN, 0.1% FA) over 90min before ramping to 37% buffer B (10min), ramp to 80% buffer B (10min) and sustained for 10min (total separation method time 120min). For the 50 samples per day method, samples were separated at 400nl/min using a linear gradient from 5% to 30% buffer B (99.9% ACN, 0.1% FA) over 16.8min before ramping to 95% buffer B (0.5min) and sustained for 2.4min (total separation method time 19.7min).

### 180 samples per day UHPLC settings

Samples were analyzed on a M-class (Waters, USA) coupled to a timsTOF Pro (Bruker) equipped with a CaptiveSpray source. Peptides were separated on a 5cm X 150μm analytical column, 1.6μm C18 beads with a packed emitter tip (IonOpticks, Australia) using a constant flow rate of 2μl/min. The column was maintained at room temperature. Sample was injected into a sample loop which takes approximately 0.5min. Mobile phase at 100% buffer A continues to flow over the analytical column during this period facilitating column equilibration. The sample loop was switched on-line for 1min at 100% buffer A. A linear gradient begins at 1.2min from 5% to 34% buffer B over 5min before ramping to 80% buffer B (0.5min) and sustained for 0.3min. Mobile phase is then ramped back to 100% buffer A (0.2min) and sustained for 0.3min (this period also contributes to column equilibration) (total method time 7.5min + 0.5min injection).

### timsTOF Pro settings

The timsTOF Pro (Bruker) was operated in PASEF mode using Compass Hystar 5.0.36.0. Settings for the 11 samples per day method were as follows: Mass Range 100 to 1700m/z, 1/K0 Start 0.6 V.·/cm^2^ End 1.6 V·s/cm^2^, Ramp time 110.1ms, Lock Duty Cycle to 100%, Capillary Voltage 1600V, Dry Gas 3 l/min, Dry Temp 180°C, PASEF settings: 10 MS/MS scans (total cycle time 1.27sec), charge range 0-5, active exclusion for 0.4 min, Scheduling Target intensity 10000, Intensity threshold 2500, CID collision energy 42eV. Settings for the 50 and 180 samples per day method were as follows: Mass Range 100 to 1700m/z, 1/K0 Start 0.85 V·s/cm^2^ End 1.3 V·s/cm^2^, Ramp time 100ms, Lock Duty Cycle to 100%, Capillary Voltage 1600V, Dry Gas 3 l/min, Dry Temp 180°C, PASEF settings: 4 MS/MS scans (total cycle time 0.53sec), charge range 0-5, active exclusion for 0.4 min, Scheduling Target intensity 24000, Intensity threshold 2000, CID collision energy 42eV.

### Raw data processing and analysis

All raw files were analyzed by MaxQuant (v1.6.10.43 or 1.6.14) software using the integrated Andromeda search engine. Experiment type was set as TIMS-DDA with no modification to default settings. Data was searched against the human Uniprot Reference Proteome with isoforms (downloaded March 2019) and a separate reverse decoy database using a strict trypsin specificity allowing up to 2 missed cleavages. The minimum required peptide length was set to 7 amino acids. Modifications: Carbamidomethylation of Cys was set as a fixed modification, while N-acetylation of proteins and oxidation of Met were set as variable modifications. First search peptide tolerance was set at 20ppm and main search set at 6ppm (other settings left as default). Single shot samples and fractions were assigned as separate parameter groups, fractions were not assigned an experiment name or fraction number and matching between runs was turned on. Maximum peptide mass [Da] was set at 8000.

## Results

Traditionally, single injection shotgun proteomics workflows have utilized liquid chromatography gradients of >60min to ensure that chromatographic peaks are wide enough to allow for multiple MS1 measurements across the peaks facilitating accurate label-free quantitation. In data-dependent acquisition modes, the minimum duty cycle of the instrument dictates the required peak widths. With the advent of the timsTOF Pro mass spectrometer and PASEF acquisition modes^7^ we now have the ability to perform data-dependent acquisition using sub-one second duty cycles allowing for significantly reduced peak widths and shorter LC gradients. Shotgun proteomic workflows designed for >60min gradients achieve high numbers of peptide identifications and robust quantitation but limit the number of samples analyzed to around 10-12 per day (Fig. 1A, Supp. Fig. 1). To maximize the efficiency of peptide and protein identification we took advantage of the reduced duty cycle time on the timsTOF Pro with PASEF by designing a series of short gradients that were paired with analytical columns of specific dimensions to optimize results from the reduced gradient lengths (Fig. 1B and C). We first utilized an analytical column configuration measuring 15cm X 75μm with 1.6um C18 particles packed into the emitter coupled to ultra-high-performance liquid chromatography (UHPLC) instrument capable of loading and equilibrating analytical columns under set backpressure conditions. Equilibrating and sample loading the analytical column using 980 bar backpressure allows for the combined steps to be reduced to 6 minutes for a 1μl sample injection. These UHPLC systems require a period of valve switching and gradient preparation between the equilibration/load steps and the sample gradient introducing a period of around 2.8 minutes of “dead time”. A 17min sample gradient is then performed with an additional 3 minutes to allow for ramping and washing the column in a high concentration of organic mobile phase. Including sample pick-up, this workflow results in a sample through-put of approximately 50 samples per day (Fig. 1B, Supp. Fig. 2). Next, we tested a shorter column measuring 5cm X 150μm with 1.6um C18 particles packed into the emitter tip coupled to a UHPLC that retains a stable flowrate of 2μl/min and loads and injects samples by using an injection valve to add or remove the sample loop from the flow path. In this arrangement, the equilibration and sample injection steps are performed concurrently as the mobile phase continues to flow over the analytical column during a 1μl sample injection. The sample loop is then switched online, with the 1μl of sample and an additional 1μl of mobile phase passing through the loop in 1min before switching the loop off-line. A 5min gradient is then performed with an additional minute to allow for ramping and washing the column in a high concentration of organic mobile phase. The total run time including sample injection is 8min equating to 180 samples per day (Fig. 1C, Supp. Fig. 3).

**Figure 1.**
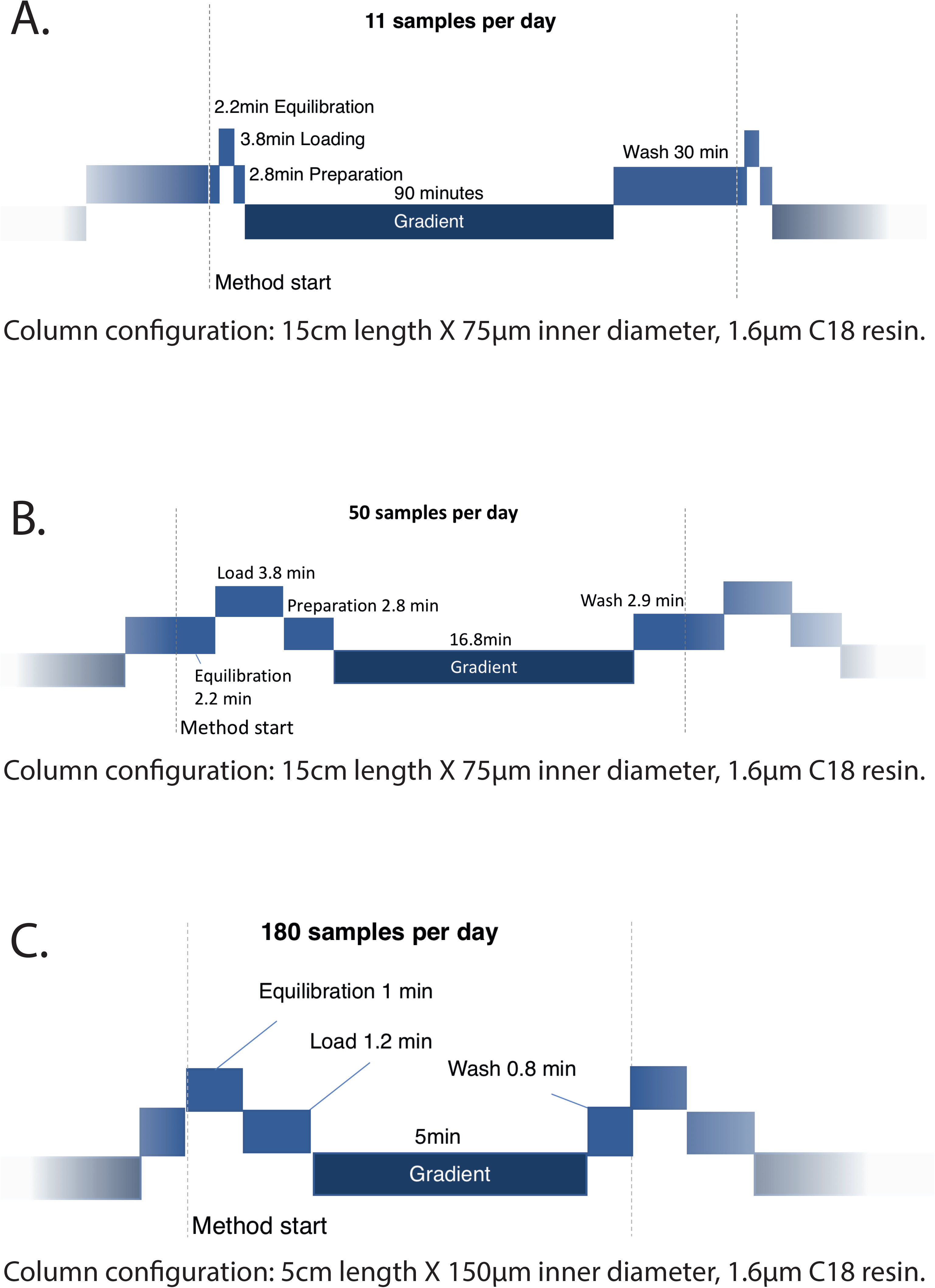
Schematics of UHPLC methods. A) 90min sample gradient (11 samples per day). Total method time 128.8min. B) 16.8min sample gradient (50 samples per day). Total method time 28.5min. C) 5min sample gradient (180 samples per day). Total method time 8min.

To determine the outcome of utilizing a 17min gradient (50 samples per day) method to increase sample throughput we performed 50 injections of 200ng Hela tryptic digest in addition to running 4 technical replicates of a Hela tryptic digest that was separated into 12 fractions using high pH reversed phase fractionation in a stage-tip format. Comparing the 50 single shot samples to the peptides observed in the 12 fractions and employing the ‘matching between runs’ feature in MaxQuant, we identified an average of 30,329 unique peptide sequences (cumulative total of 51,503) (Fig. 2A) and 5,931 proteins (cumulative total 7,244)(Fig. 2B) in each single shot replicate. When we analyzed cumulative peptide and protein identifications across all fractions, we observed a median of 47,918 unique peptide sequences (cumulative total of 57,212)(Fig. 2C) and 7,260 proteins (cumulative total of 7,573)(Fig. 2D). We repeated this process using our 5min gradient (180 samples per day) by performing 50 injections of Hela tryptic digest in addition to running 2 technical replicate of 12 reversed phase high pH fractions of a Hela tryptic digest to match to. From these samples we observe an average of 16,538 unique peptide sequences (cumulative total of 28,250)(Fig. 2A) and 3,666 proteins (cumulative total of 4,659)(Fig. 2B) in each single shot replicate. When we analyzed the cumulative peptide and protein identifications across all fractions, we observed 31,637 unique peptide sequences (Fig. 2C) and 4,946 proteins (Fig. 2D). Together, these results demonstrate that by utilizing fractions to create a peptide library to match identifications to, we can achieve high numbers of peptide and protein identifications using short sample gradients.

**Figure 2.**
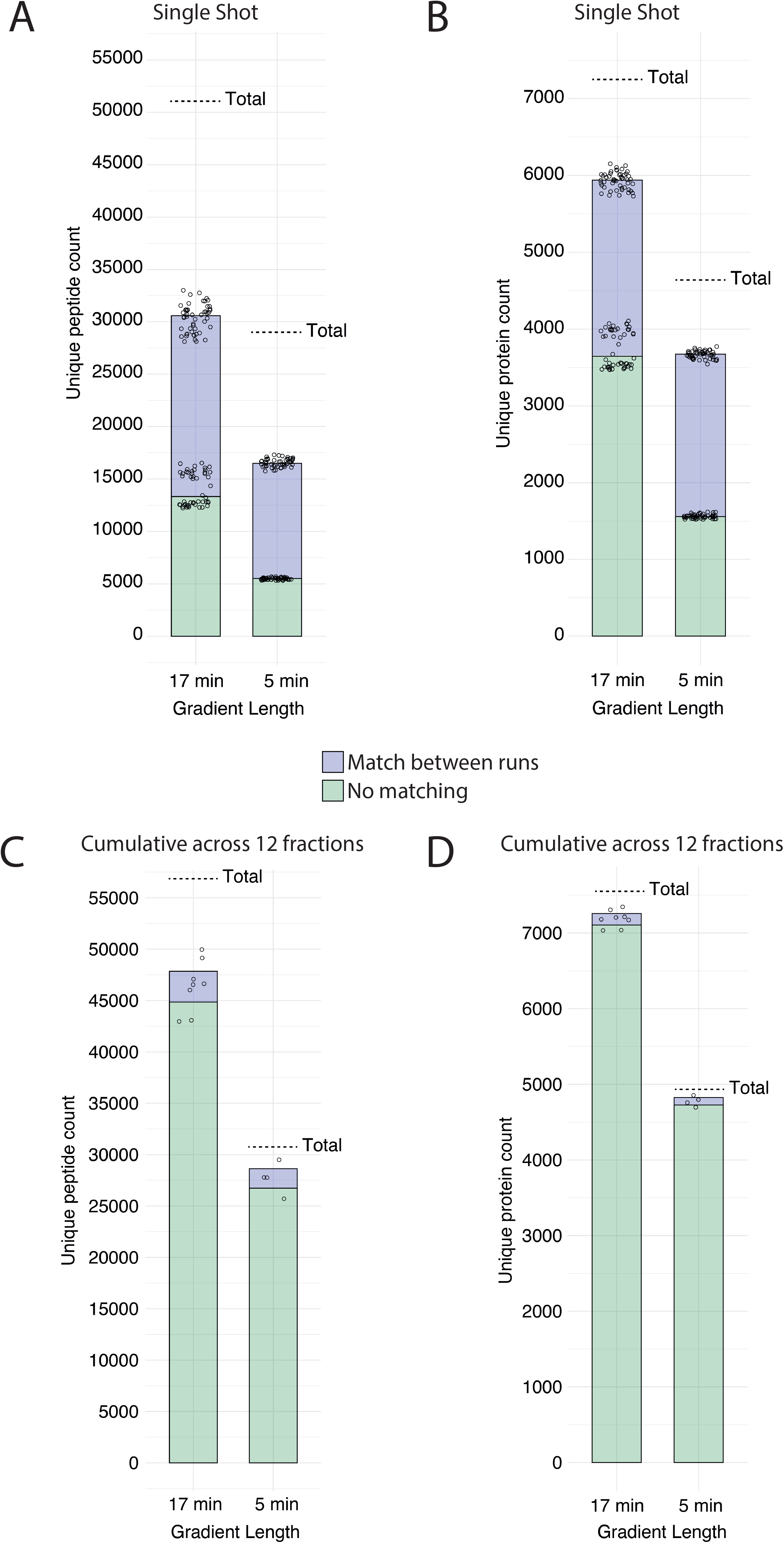
High numbers of peptide and protein identifications achieved from short gradients. A and B) Unique peptide sequence and unique protein group identifications from Hela tryptic digest using 17min (200ng injection, n= 50) and 5min (80ng injection, n=50) gradient single shot runs. Single shot samples were matched to 12 high pH reversed phase fractions of the same samples. C and D) Cumulative total of 12 fractions used in A and B. 17min gradient, n = 4; 5min gradient, n = 2. All dots represent individual replicate values, bars represent the average value across replicates and dashed line indicates the cumulative total identifications across all runs. Colors distinguish the identification with the Match between runs feature enabled in MaxQuant or not.

We next determined the effect of reduced gradient lengths and column configurations on peak characteristics. The median full width at half maximum (FWHM) of peaks on a 90min gradient was 10.1 seconds over three replicates. Shorter gradient lengths led to a reduction in the median FWHM peak with to 3.7 and 2.2 seconds for the 17min gradient and 5min gradients, respectively (Fig. 3A). Overlaying the chromatographic traces from multiple replicates demonstrated high inter-replicate reproducibility with very little variation observed between runs for both the 17min gradient (Fig. 3B) and the 5min gradient (Fig. 3C). To demonstrate robust performance over a larger sample set we mapped the retention time for multiple peptides from different protein groups across all replicates. Results from the 17min and 5min gradients demonstrated very little divergence in retention time across the complete datasets (Fig. 4A and B).

**Figure 3.**
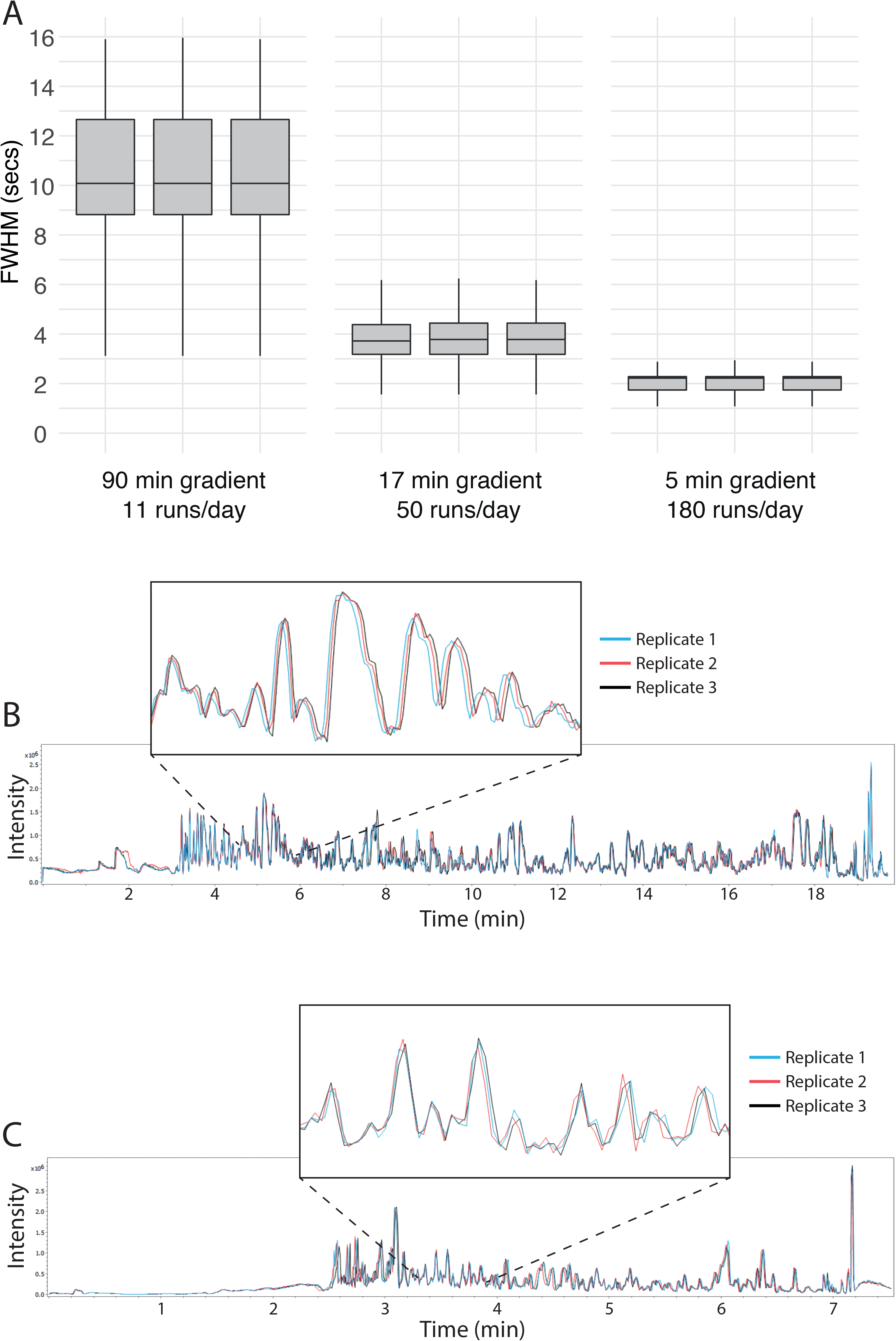
Chromatographic peak characteristics from short gradient methods. A) Box plot of identified peptides full width at half maximum (FWHM) in seconds for column length and gradient duration combination. Three replicates of 200ng Hela tryptic digest injections shown. Line indicates median FWHM for each sample. Outliers were omitted from the plot. B and C) Comparison of three base peak chromatograms from a HeLa tryptic digest (200ng injection) for the 17 (B) and 5 (C) minute gradients.

**Figure 4.**
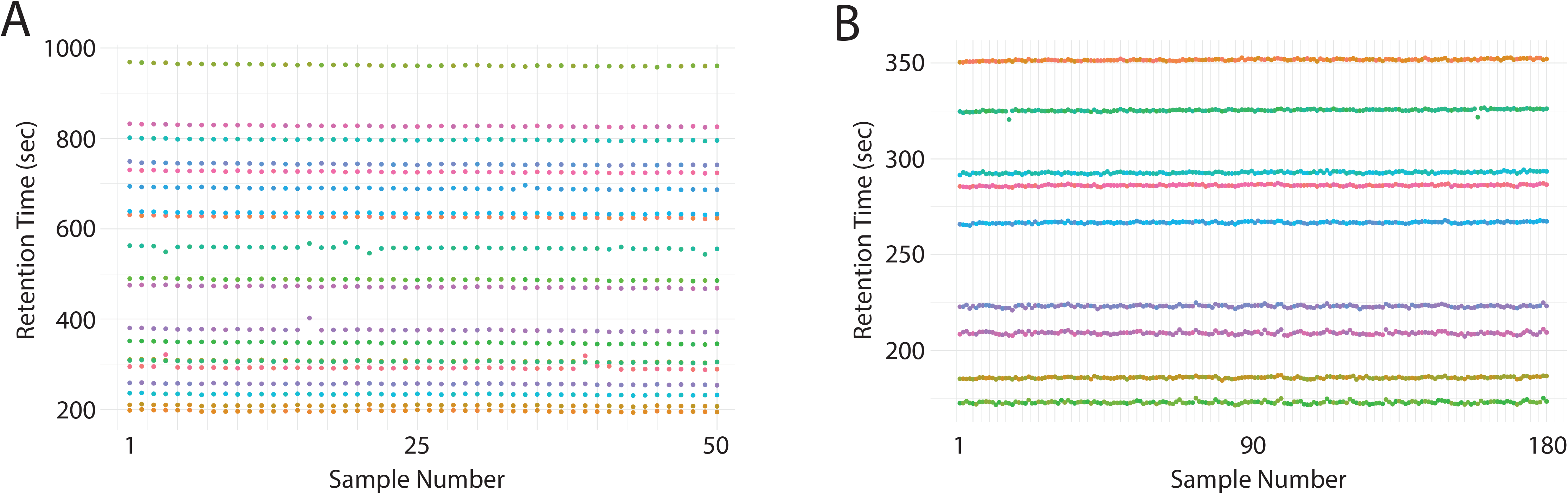
Short gradient methods allow reproducible analysis of samples. A and B) Retention time stability of selected peptides from 200ng (17min, 50 samples) or 80ng (5min, 180 samples) injections of a Hela tryptic digest that were identified across all samples. Each peptide is represented by a different color.

Having demonstrated an ability to achieve high numbers of identifications in a common standard laboratory sample, we next sought to analyze a sample with clinical relevance. Blood plasma is widely used for clinical diagnostics and is a rich source of potential biomarkers across a broad range of diseases. A single sample of Plasma was collected, processed and tryptically digested before being injected 180 times using the 5min gradient method. The same sample was also separated into 2 technical replicates of 12 fractions using high pH revered phase fractionation in a stage-tip format to generate a peptide library. This analysis successfully identified an average of 1,694 unique peptide sequences (cumulative total 2,373)(Supp. Fig. 4) and 238 proteins (cumulative total 293)(Fig. 5A) per sample. Analysis of protein intensities across the 180 runs demonstrated a high degree of correlation between the samples with an average correlation of 0.975 (Fig. 5B). These results demonstrate that the high-throughput method can be utilized to characterize clinically relevant samples and achieve a large number of protein identifications with high reproducibility.

**Figure 5.**
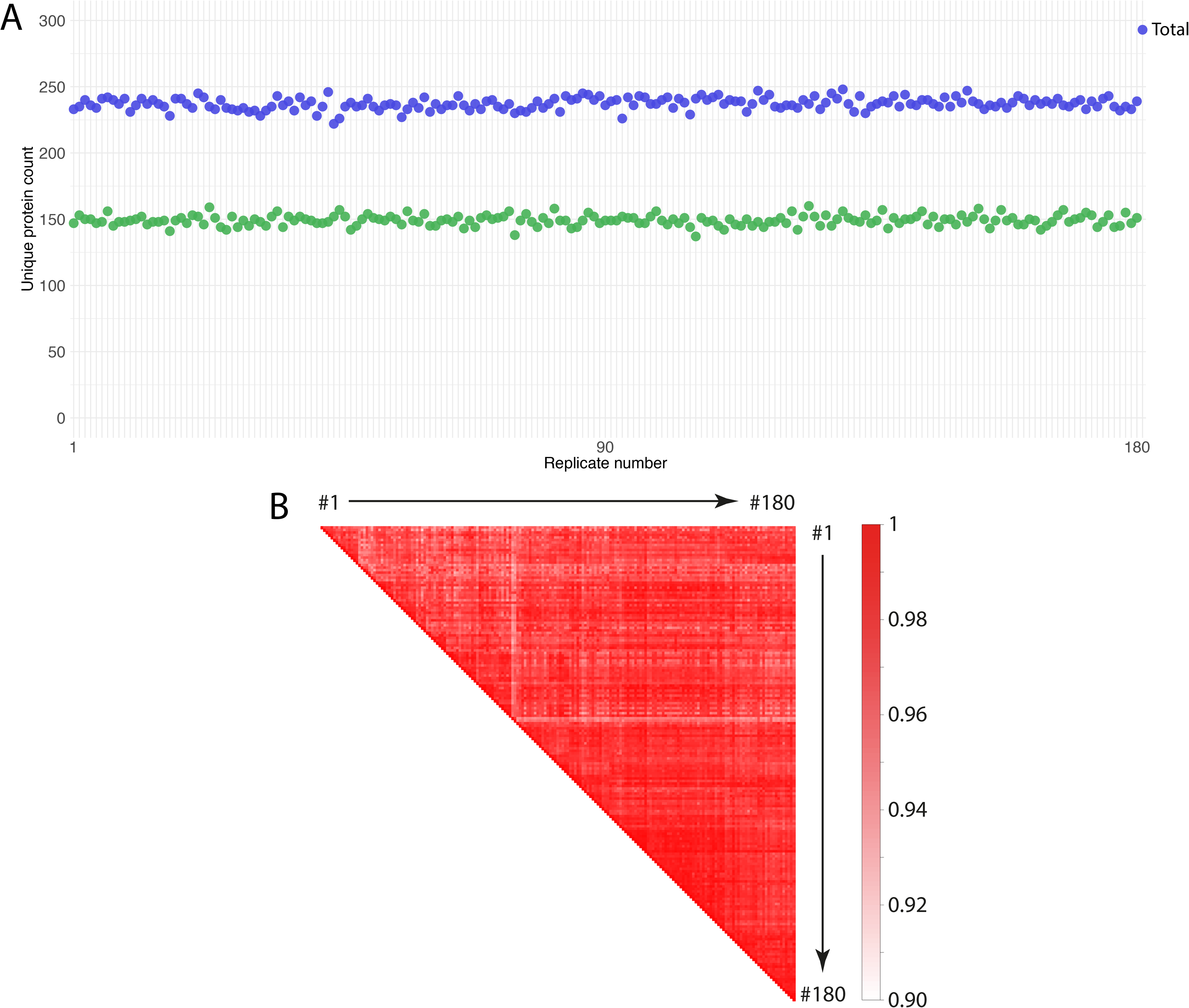
Short gradients facilitate analysis of Plasma samples. A) Unique protein group identifications from Plasma digest using 180 (5min gradient; 50ng injection, n= 180) samples per day single shot runs. Single shot samples were matched to 12 high pH reversed phase fractions of the same sample. B) Pearson correlation matrix comparing protein intensity measurements of all 180 plasma runs to each other. Average correlation was 0.975.

## Discussion

A key limitation to DDA based shotgun proteomics is the combination of stochastic feature sampling and an ability to generate MS2 spectra intense enough to confidently make the identification (effective sequencing speed), generally causing large numbers of missing values across sample sets. Accurate matching from a library of identified MS1 features (‘Accurate Mass Tagging’ and ‘match between runs’^11,12^ has widely been implemented over the last decade even though stringent control of false discovery rates has yet to be fully implemented. Here we demonstrate the effectiveness of combining high peak capacity chromatography on short gradients with the timsTOFs speed and CCS enhanced selectivity to provide an extremely high yielding proteomics discovery workflow.

The high speed and sensitivity of the timsTOF Pro utilizing the PASEF mode of operation has been clearly demonstrated previously^7^. However, workflows that fully take advantage of the reduction in cycle times, increased sequencing speed and CCS assisted matching has yet to ascertained. Here we evaluate the ability of the PASEF shotgun proteomics workflow, utilizing rapid gradients on high peak capacity packed emitter columns. By combining these short gradients with the recently implemented CCS enhanced library matching in MaxQuant^9^ with stage-tip high pH reversed phase fractionation, we demonstrate very high identification rates for short gradients with complex peptide mixtures. Thus, the combination of high pH library generation, additional feature matching selectivity using CCS values and high peak capacity chromatography provides a deep and very high through-put label free quantification workflow. Interestingly, the level of proteins matched across the individual samples nears the level of proteins found across the 12 stage-tip fractions, suggesting that increasing the number of off-line high pH fractions could potentially improve the performance of shotgun matching depth even further. Further, the discrepancy of unique peptide counts between the individual technical replicates and the cumulative total count suggests that improvements to MS1 acquisition parameters and more advanced matching strategies that incorporate FDR based matched cut-offs may also further improve the data completeness. The rapid acquisition of 12 high pH fractions in 6 hours (for 50 samples per day method) enables routine high pH fractionation for each experiment. This depth of sequencing, speed and reported instrument robustness^7^ now provides a very appealing setup for clinical analysis, where historically, numbers of patient samples have been limited due to the low through-put nature of long gradient LC-MS analysis, generally limiting the quality and reproducibility of the results^13^. Higher throughput and larger numbers of patient samples will undoubtedly result in greater robustness of potential identified markers and has the additional benefit of more reliable investigation of experimental and patient confounders^14^.

The performance obtained here is accessible as it uses a defined series of available methods, analysis using freely available software and commercially available columns. The methods described here do not require a specialized skill set and are simple to implement making this accessible to a broad range of proteomics research laboratories. Challenges that remain include software analysis time (which needs to be improved by a factor of 2-3 to allow laboratories to operate inside of the dreaded time-debt scenario, where the time it takes to feature detect and search the data takes longer than the acquisition time). The ultra-fast acquisition of proteomes in the context of known physical limitations of MS dynamic range strongly suggests future strategies for deep proteome analysis will likely involve high pH reversed phase fractionation and even more rapid acquisition modes.

## Acknowledgements

This work was supported by operational infrastructure grants through the Australian Government IRISS and the Victorian State Government OIS 9000220.

## Figure legends

Supplementary Figure 1.

Schematic of a single 11 samples per day (90min gradient) method. A) Trace of Pump A pressure during run. B) Trace of percentage of buffer B during run. C) Total Ion Chromatogram plot across gradient. D) Base Peak Chromatogram across gradient.

Supplementary Figure 2.

Schematic of a single 50 samples per day (17min gradient) method. A) Trace of Pump A pressure during run. B) Trace of percentage of buffer B during run. C) Total Ion Chromatogram plot across gradient. D) Base Peak Chromatogram across gradient.

Supplementary Figure 3.

Schematic of 15 X 180 samples per day (5min gradient) method. A) Trace of Pump A pressure during runs. B) Trace of percentage of buffer B during runs. C) Total Ion Chromatogram plot across gradients. D) Base Peak Chromatogram across gradients.

Supplementary Figure 4.

Unique peptide sequence identifications from Plasma digest using 180 (5min gradient; 50ng injection, n= 180) samples per day single shot runs. Single shot samples were matched to 12 high pH reversed phase fractions of the same samples.

