## Supplementary figures and images for "Simplified high-throughput methods for deep proteome analysis on the timsTOF Pro"

### Supplemental Figure 1

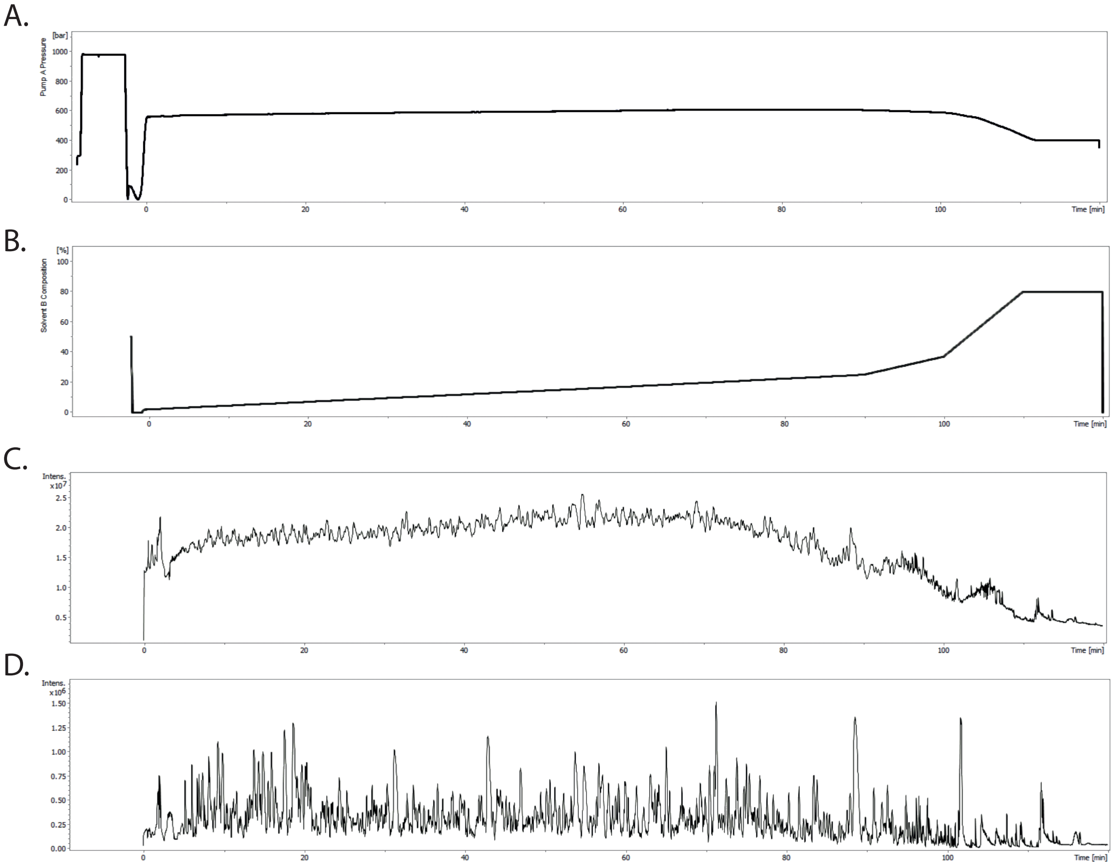

### Supplemental Figure 2

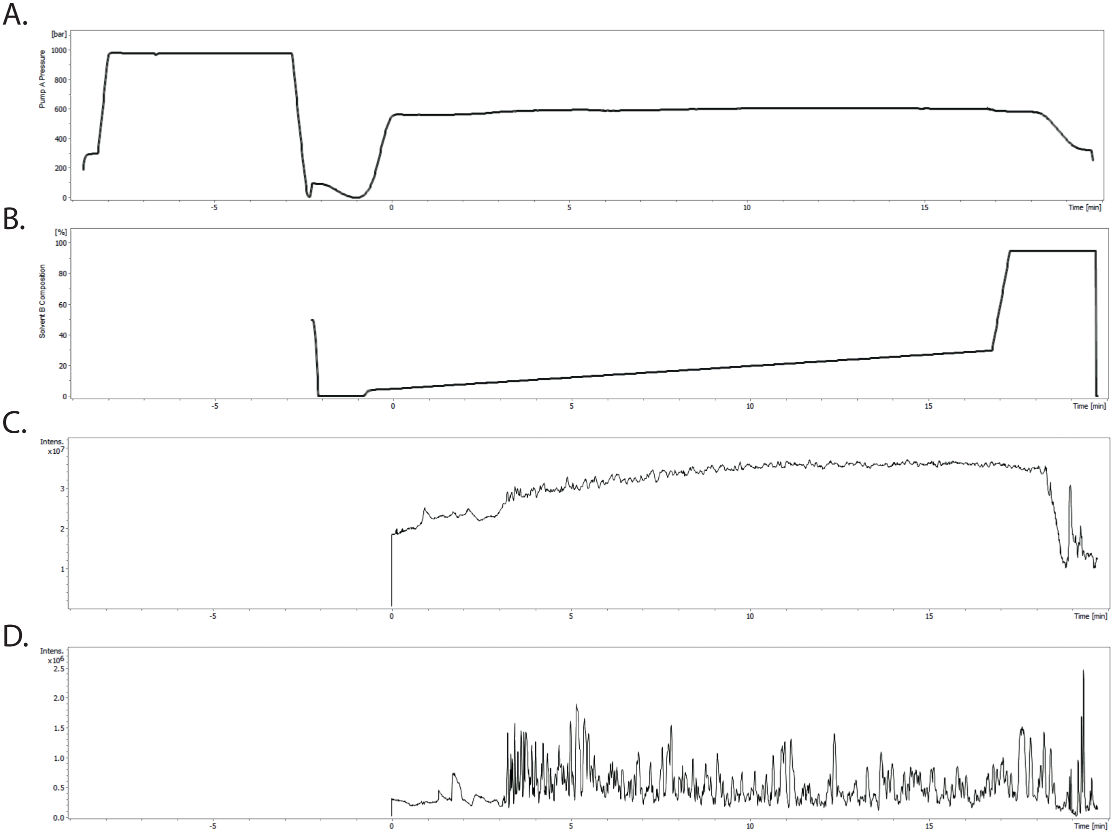

### Supplemental Figure 3

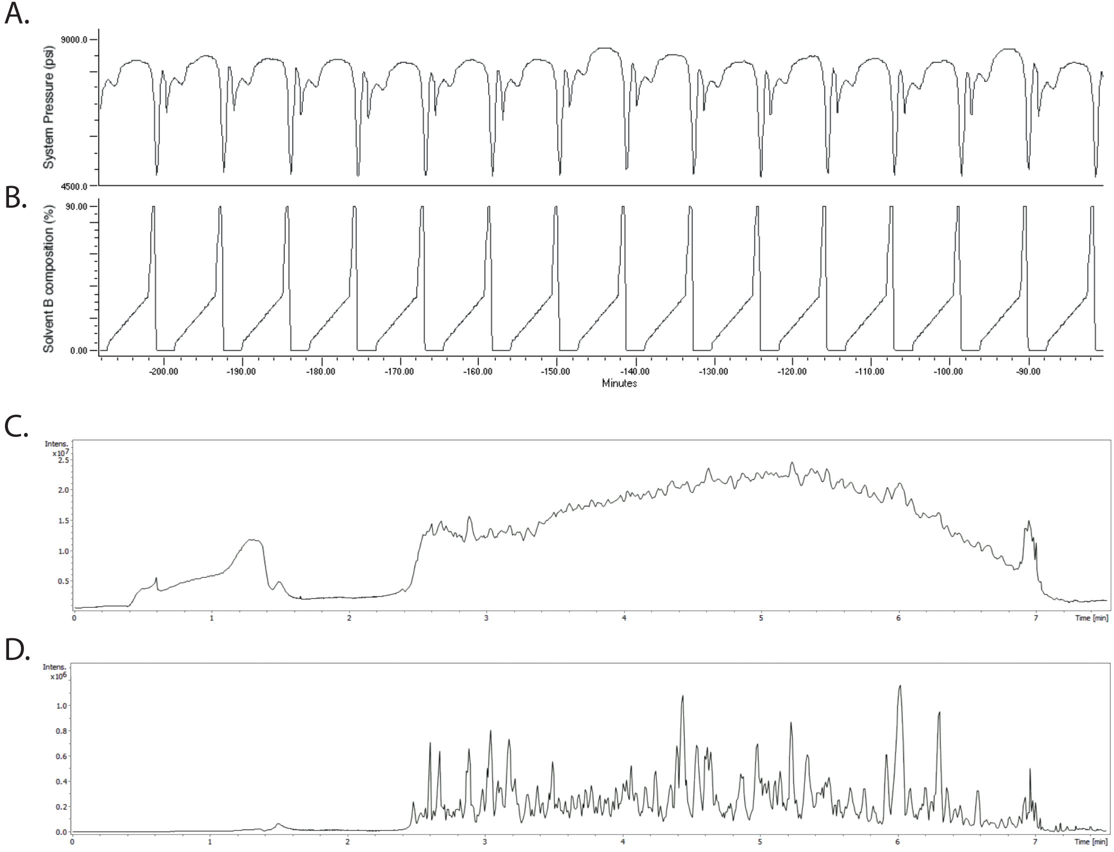

### Supplemental Figure 4

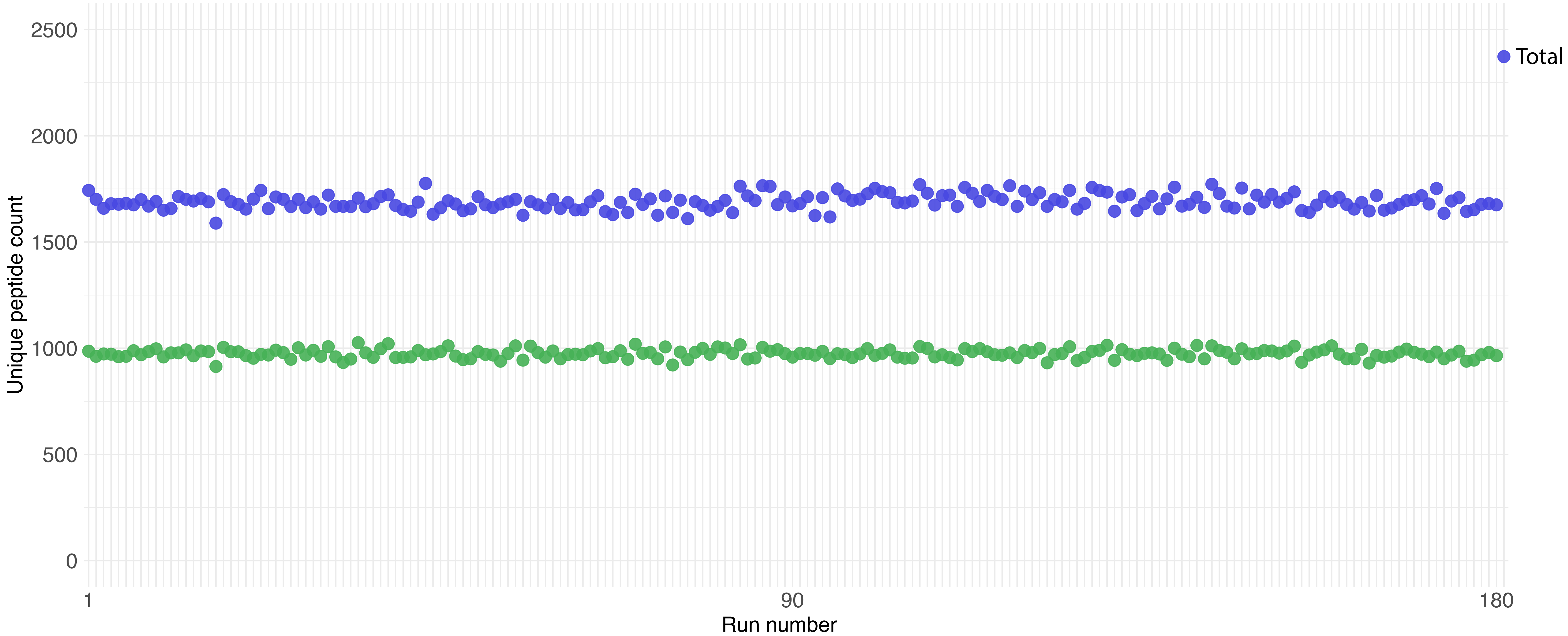
